# The RNA sensor MDA5 detects SARS-CoV-2 infection

**DOI:** 10.1101/2021.03.26.437180

**Authors:** Natalia G. Sampaio, Lise Chauveau, Jonny Hertzog, Anne Bridgeman, Gerissa Fowler, Jurgen P. Moonen, Maeva Dupont, Rebecca A. Russell, Marko Noerenberg, Jan Rehwinkel

## Abstract

Human cells respond to infection by SARS-CoV-2, the virus that causes COVID-19, by producing cytokines including type I and III interferons (IFNs) and proinflammatory factors such as IL6 and TNF. IFNs can limit SARS-CoV-2 replication but cytokine imbalance contributes to severe COVID-19. We studied how cells detect SARS-CoV-2 infection. We report that the cytosolic RNA sensor MDA5 was required for type I and III IFN induction in the lung cancer cell line Calu-3 upon SARS-CoV-2 infection. Type I and III IFN induction further required MAVS and IRF3. In contrast, induction of IL6 and TNF was independent of the MDA5-MAVS-IRF3 axis in this setting. We further found that SARS-CoV-2 infection inhibited the ability of cells to respond to IFNs. In sum, we identified MDA5 as a cellular sensor for SARS-CoV-2 infection that induced type I and III IFNs.

## Introduction

Severe acute respiratory syndrome coronavirus 2 (SARS-CoV-2) emerged at the end of 2019 and is causing an ongoing global pandemic. As of March 26^th^, 2021, it has infected 124,535,520 and killed 2,738,876 patients worldwide (https://covid19.who.int/). SARS-CoV-2 causes Coronavirus Disease 2019 (COVID-19), a respiratory disease that in some patients results in severe pneumonia and acute respiratory distress syndrome leading to death. Severe disease is linked to exacerbated inflammation with increased production of pro-inflammatory cytokines such as TNF and IL-6, and delayed type I and type III interferon (IFN) responses^1–4^. Although serum levels of type I and III IFNs are low or undetectable in many patients, the increased expression of genes known to be induced by IFN (called interferon stimulated genes; ISGs) suggests that production of IFN occurs. The importance of the IFN system in controlling disease was confirmed by the discovery of an association between severe disease and inborn errors in IFN immunity as well as autoantibodies against type I IFNs^2,5^. Moreover, recent studies show that intranasal, but not intravenous, administration of type I IFN improves disease outcome in an *in vivo* hamster model as well as in a phase 2 clinical trial, supporting a crucial role for the type I IFN response in COVID-19^6–8^. Similarly, pegylated IFNλ1, a type III IFN, shows prophylactic and therapeutic benefits in a mouse model of SARS-CoV-2, and is currently being tested in clinical trials^9,10^. The main replication sites of SARS-CoV-2 in patients are the upper and lower respiratory tract, where the virus infects airway and alveolar epithelial cells, vascular endothelial cells, and alveolar macrophages^11^. Uncovering how these cells detect SARS-CoV-2 infection to produce type I and III IFNs and other pro-inflammatory cytokines is therefore of great importance to understanding the pathology of COVID-19.

SARS-CoV-2 is a member of the beta-coronavirus family and is closely related to other viruses that have caused outbreaks in the last two decades: SARS-CoV in 2003 and MERS-CoV in 2013. Its genome is a ~30kb positive-sense single-stranded RNA that shares ~80% sequence identity with SARS-CoV and ~50% sequence identity with MERS-CoV^11^. Nucleic acid sensors mediate the early detection and host response to virus infections. These sensors recognise either viral nucleic acids or ‘unusual’ cellular nucleic acids present upon infection^12^. Cytosolic nucleic acid sensors from the RIG-I-Like Receptor (RLR) family have been identified as important PRRs that sense coronaviruses^9,13^. The two signalling receptors in this family are retinoic acid-inducible gene I (RIG-I) and melanoma differentiation-associated protein 5 (MDA5), which detect RNAs with specific structures such as 5’-triphosphate or 5’-diphosphate ends^14,15^. Once activated, RIG-I and MDA5 interact with the adaptor mitochondrial antiviral-signalling protein (MAVS), triggering a signalling cascade involving TANK-binding kinase 1 (TBK1) and interferon regulatory factor 3 (IRF3) that ultimately induce the expression of type I IFNs (including IFNα and IFNβ), type III IFNs (also known as IFNλ) and other antiviral genes. Secreted type I and III IFNs signal in a paracrine and autocrine manner through their receptors IFNAR and IFNLR, respectively. Both receptors activate the JAK/STAT signalling pathway, leading to the expression of ISGs, some of which encode anti-viral proteins such as MxA^16^. In some infections with RNA viruses such as Dengue virus, the cytosolic DNA sensing pathway involving cGAMP synthase (cGAS) and stimulator of interferon genes (STING) detects mitochondrial DNA leaked from damaged mitochondria and thereby contributes to IFN induction^17,18^. Whether this also happens during coronavirus infection is unknown.

In this study, we identified Calu-3 cells as a suitable model to study innate immune responses to SARS-CoV-2 infection. Using shRNA depletion and CRISPR-Cas9 knockout, we found that MDA5 was an important cytosolic sensor for SARS-CoV-2 in these cells. The MDA5-MAVS-IRF3 signalling axis was necessary for production of type I and III IFNs, but not pro-inflammatory cytokines, in response to SARS-CoV-2 infection. We further show that expression of the ISG MxA is mainly induced in bystander non-infected cells in this system.

## Results

In order to identify innate immune sensors that detect SARS-CoV-2, we screened a number of cell lines for permissiveness to infection and immune response (Fig 1A). Calu-3, HEK293 and Huh7 cells were permissive to infection, with expression of SARS-CoV-2 nucleocapsid (N) RNA detectable in these cells 24 hours after infection (Fig 1B). However, only Calu-3 cells showed both visible signs of infection by microscopy (Fig 1A) and an innate immune response to the virus (Fig 1B). Upon SARS-CoV-2 infection, Calu-3 cells upregulated expression of type I and III IFNs (*IFNB1, IFNL1*), pro-inflammatory cytokines (*IL6, TNF*) and ISGs (*IFIT1, IFIH1, MX1*). Calu-3 are adenocarcinoma-derived lung epithelial cells and as such are related to one of the cell types infected by SARS-CoV-2 in the respiratory tract. In addition, Calu-3 cells expressed many of the proteins required for the major signalling pathways activated during viral infection: cGAS and STING (DNA sensing); RIG-I, MDA5 and MAVS (RNA sensing); the TLR adaptor MyD88; the kinase TBK1; and the transcription factors IRF3 and STAT1/2 (Fig 1C). These cells further responded to type I IFN stimulation by phosphorylation of STAT1 and STAT2 (Fig 1C). Together, these data established that Calu-3 cells were a good model for investigating innate immune responses to SARS-CoV-2 infection.

**Figure 1.**
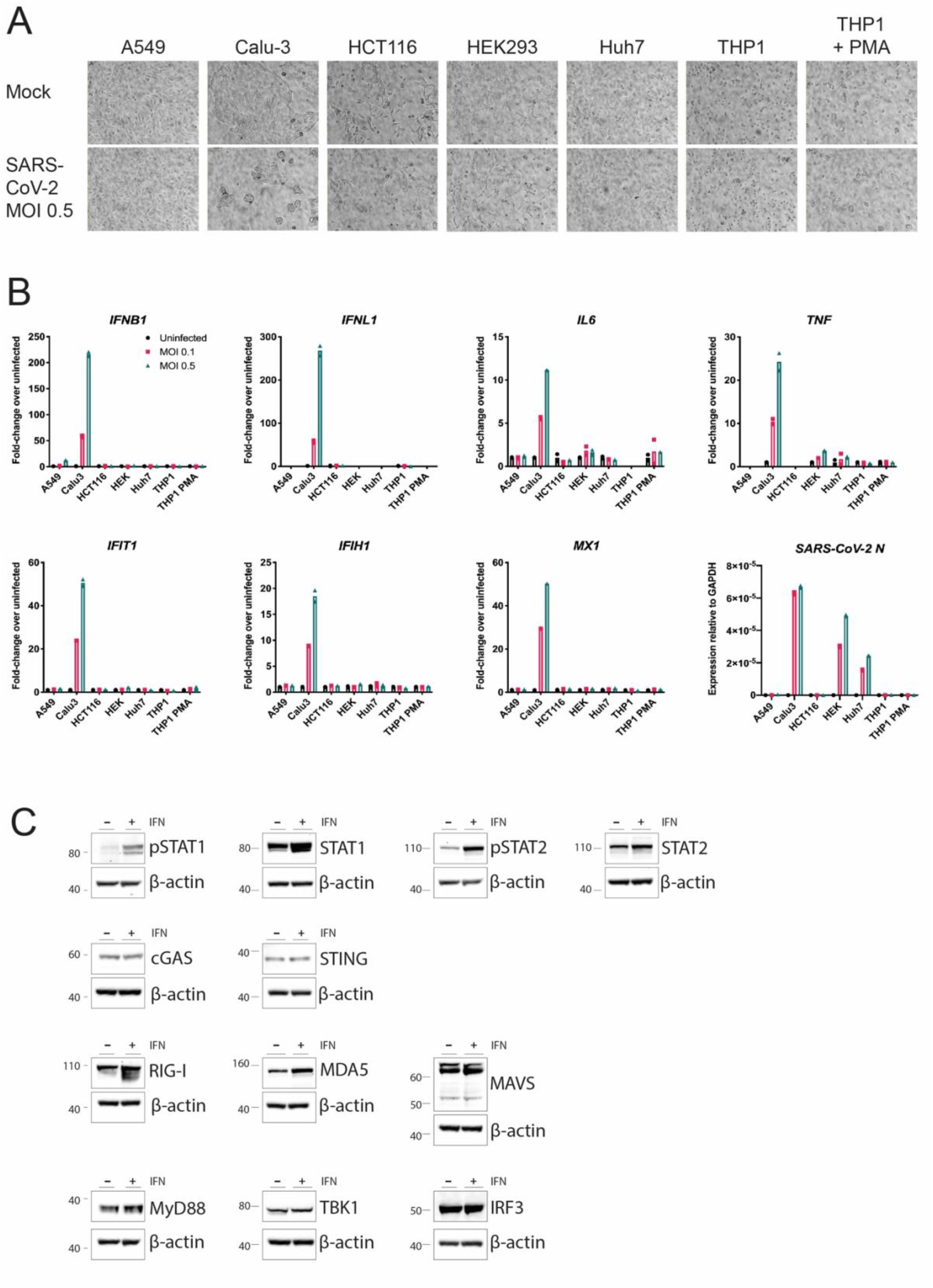
Calu-3 cells respond to SARS-CoV-2 infection by upregulating type I and III IFNs, ISGs and cytokines. (A, B) The indicated cell lines were mock-infected or infected with SARS-CoV-2 (MOI=0.1 or 0.5) for 24 hours prior to brightfield imaging (A) or RNA extraction and RT-qPCR for the indicated transcripts (B). Data in (B) are relative to *GAPDH* expression. (C) Calu-3 cells were stimulated with 100 U/ml IFN-A/D for 16 hours. Cell lysates were analysed by western blot using the indicated antibodies. Data are from a single experiment. Data points in (B) are from technical duplicates and bars indicate the average.

To investigate if RIG-I-like receptors (RLRs) were involved in virus sensing, we first utilised a lentiviral shRNA knockdown approach targeting MAVS, the downstream signalling adaptor for both RIG-I and MDA5. Two independent shRNAs were tested: both reduced MAVS protein and mRNA levels, with shRNA-MAVS-45 having the more potent effect (Fig 2A,B). We therefore chose shRNA-MAVS-45 for further experiments and infected Calu-3 cells transduced with this shRNA with SARS-CoV-2 using a multiplicity of infection (MOI) of 0.1. 48 hours after infection, RNA was extracted from cells and viral and cellular RNAs were analysed by RT-qPCR. The levels of SARS-CoV-2-N RNA were slightly increased in MAVS-depleted cells (Fig 2C). Importantly, induction of type I and III IFN mRNAs and ISGs in response to SARS-CoV-2 was reduced in cells depleted of MAVS (Fig 2D). However, expression of the mRNAs encoding the pro-inflammatory cytokines TNF and IL6 was not affected by knockdown of MAVS (Fig 2D). Taken together, these data suggest that RLRs are necessary for the IFN response to SARS-CoV-2 in Calu-3 cells.

**Figure 2.**
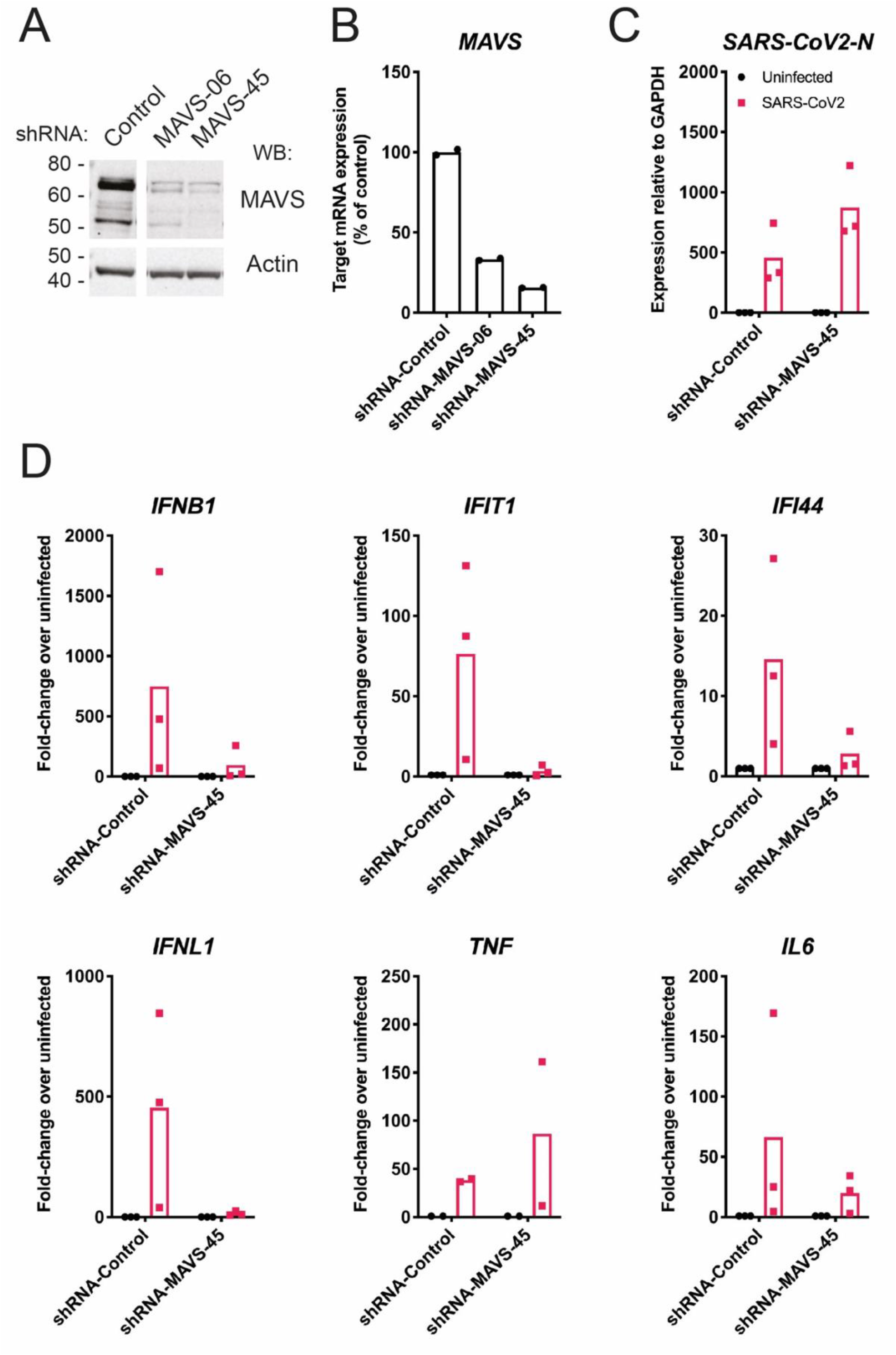
MAVS is required for the type I and III IFN response to SARS-CoV-2. (A, B) Calu-3 cells were depleted of MAVS by lentiviral shRNA delivery using two independent shRNAs (shRNA-MAVS-06 and shRNA-MAVS-45). Knockdown efficiency was assessed by western blot (A) and RT-qPCR (B). The control shRNA targeted GFP, which is absent in Calu-3 cells. (C, D) Calu-3 cells depleted of MAVS using shRNA-MAVS-45 were infected with SARS-CoV-2 (MOI=0.1) for 48 hours, followed by RNA extraction and RT-qPCR for the indicated transcripts. Data are relative to *GAPDH* expression. Data in (A, B) are representative of two independent biological repeats. Data in (C-D) are pooled from 2-3 independent biological repeats, with bars representing the average.

Next, we studied the roles of the two signalling RLRs, MDA5 and RIG-I, in the sensing of SARS-CoV-2 using a knock-out approach. We found that Calu-3 cells were not amenable to selection of clones from single cells and therefore opted to use a lenti-CRISPR approach. We transduced Calu-3 cells with lentiviruses expressing Cas9, an sgRNA and the puromycin resistance gene. Using puromycin selection, we generated polyclonal cell lines. We included sgRNAs targeting MAVS, MDA5 and RIG-I. To determine if the cytosolic DNA sensing pathway was activated during SARS-CoV-2 infection, we also targeted STING. IRF3 is activated downstream of both STING and MAVS; therefore, we included a IRF3 sgRNA. Protein levels of MDA5, RIG-I, STING and IRF3 were notably reduced by this approach, whereas the targeting of MAVS was less efficient (Fig 3A). Next, we infected these cells with SARS-CoV-2 (MOI=0.1) and analysed their response by RT-qPCR after 48 hours. Cells lacking MDA5 or IRF3 showed minimal induction of type I IFN mRNAs and ISGs in response to SARS-CoV-2 infection (Fig 3B). This effect was uncoupled from the pro-inflammatory response, as lack of MDA5 and IRF3 did not affect the expression of *TNF*, and minimally affected expression of *IL6*, in response to infection (Fig 3B). Lack of RIG-I or STING had no effect on the cellular response to SARS-CoV-2, suggesting that neither RIG-I nor the cGAS-STING pathway were involved in sensing SARS-CoV-2 in this setting. It likely that partial depletion of MAVS (Fig 3A) explains why cells transduced with MAVS sgRNA responded comparably to control cells.

**Figure 3.**
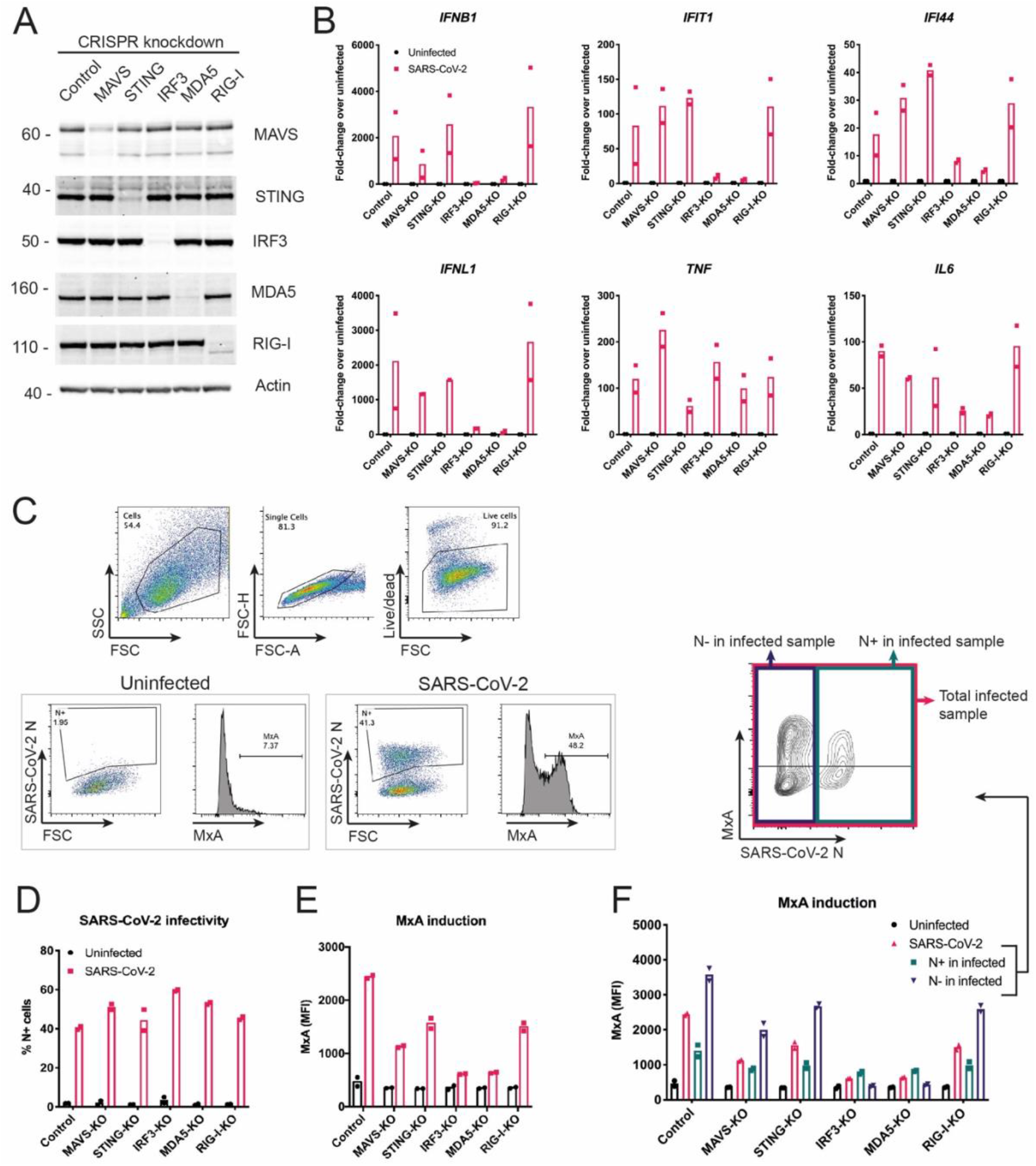
SARS-CoV-2 activates type I and III IFN responses *via* MDA5 and IRF3, and inhibits ISG induction in infected Calu-3 cells. (A) Calu-3 cells were depleted of MAVS, STING, IRF3, MDA5 or RIG-I using a lenti-CRISPR approach. Depletion of the targeted proteins was assessed by western blot. Control cells were targeted for GFP, which is absent in Calu-3 cells. (B) Cells from (A) were infected and analysed as described in Figure 2C. (C-F) Cells from (A) were infected as in Figure 2C, stained for live cells, SARS-CoV-2 N protein and MxA, and analysed by flow cytometry. Live cells were assessed for SARS-CoV-2 N protein expression (D) and MxA induction, shown as mean fluorescence intensity (MFI; E). (F) Cells in the SARS-CoV-2-infected samples were further subdivided into SARS-CoV-2 N positive (N+) and SARS-CoV-2 N negative (N-) cells, and MxA MFI was determined within these subpopulations. Data in (A, C) are representative of two independent biological repeats. Data in (B and D-F) are pooled from two independent biological repeats, with bars representing the average.

To confirm these observations with a different readout, we used intracellular staining and flow cytometry to measure protein levels of MxA, which is encoded by the ISG *MX1*, and SARS-CoV-2 N (Fig 3C). We analysed cells infected for 48 hours with SARS-CoV-2 (MOI=0.1). In this setting, ~40-60% of cells stained positive for N and there was little difference in infection levels between the knockout cells (Fig 3D). Consistent with our RT-qPCR data, targeting of MDA5 and IRF3 largely prevented upregulation of MxA in response to SARS-CoV-2 infection (Fig 3E). Targeting of STING and RIG-I partially reduced MxA induction. Intermediate effects were also observed with the inefficient MAVS sgRNA. Further exploration of the upregulation of MxA upon SARS-CoV-2-infection showed different responses in N+ and N-cells. In the uninfected N-cell population, MxA was upregulated to a greater extent compared with infected N+ cells (Fig 3F). This shows that SARS-CoV-2 infection inhibited the autocrine response of the same cell to released IFNs, whereas uninfected bystander cells were able to respond to IFNs more strongly, implicating viral antagonism of IFNAR and/or IFNLR signalling.

## Discussion

In this study, consistent with several recent reports^19–21^, we identified Calu-3 cells as a suitable model system for the study of innate immune responses to SARS-CoV-2 infection *in vitro*.These adenocarcinoma-derived lung epithelial cells expressed key proteins of RNA- and DNA-sensing pathways, responded to stimulation with type I IFN, and were highly infected by SARS-CoV-2. Importantly, SARS-CoV-2 induced transcription of the genes encoding type I and III IFNs, as well as pro-inflammatory cytokines in these cells. Of the other cell lines tested, A549, HCT116 and THP1 cells were not permissive to SARS-CoV-2 infection and did not upregulate antiviral cytokines in response to the virus. Other reports have established that A549 and THP1 cells do not support viral entry and/or replication, likely due to insufficient expression of the viral entry receptor ACE2^22,23^. Intriguingly, we could detect high levels of viral RNA but no induction of an IFN or cytokine response in HEK293 and Huh7 cells. HEK293 cells have been suggested to express sufficient ACE2 for viral entry, and allow for initial replication of the virus^24,25^. However, the infection appears to be abortive limiting production of substantial progeny virus^25^. Why do HEK293 cells fail to respond to the initial SARS-CoV-2 replication with transcriptional induction of antiviral cytokines? We have found that MDA5 protein is undetectable in HEK293 cells at baseline (data not shown) and speculate that this is the reason why SARS-CoV-2 is not adequately detected in these cells. Huh7 cells have been reported to be permissive to viral replication^4,23,26–30^. Recently, a proteomics-based approach found activation of the type I IFN system in Huh7 cells at 48 hours but not 24 hours post-infection^31^. This indicates a delayed IFN induction in Huh7 cells, potentially explaining our results obtained at the 24-hour timepoint.

We further show, using both shRNA-mediated knockdown and CRISPR/Cas9 genetic ablation, that the type I and III IFN response to SARS-CoV-2 was dependent on the RNA sensor MDA5, its downstream adapter MAVS and the transcription factor IRF3. In Calu-3 cells, RIG-I was largely dispensable for the antiviral cytokine response to infection. While our study was in preparation, other manuscripts reported MDA5 as the cellular sensor that recognises SARS-CoV-2 infection^19,20^. These reports included both siRNA-mediated knockdown and genetic ablation in Calu-3 cells. In contrast to these two studies and to our work, another study found that both MDA5 and RIG-I sense SARS-CoV-2 infection in Calu-3 cells, and that upregulation of the pro-inflammatory cytokine IL-6 was uncoupled from MDA5, but dependent on RIG-I and MAVS^21^. This study utilised an siRNA knockdown approach; whether this or other technical differences explain the disparity between these findings and what we and others report remains to be determined. Another research group reported that total RNA extracted from SARS-CoV-2-infected Vero E6 cells activated MDA5, but not RIG-I, after transfection into human lung fibroblasts^32^. Interestingly, a similar experimental approach in a different study obtained opposing results: here, type I IFN transcriptional upregulation after transfection of RNA from infected cells was dependent on RIG-I and not MDA5^33^. It would be interesting to compare side-by-side the different reporter cells used for RNA transfection by Liu et al. and Wu et al..

Intriguingly, our experiments showed that transcriptional upregulation of the inflammatory cytokines TNF and IL6 was not dependent on the MDA5-MAVS signalling axis. These cytokines are typically induced by the transcription factor NF-κB, which can be activated downstream of MAVS^34^, but is also stimulated by other PRRs, such as Toll-like receptors (TLRs)^35^. TLR3, a transmembrane receptor located in endosomes and at the cell surface, detects dsRNA and induces both an IRF3-mediated type I IFN response and an NF-κB-mediated pro-inflammatory response. Mutations in *TLR3* are associated with disease severity in patients, suggesting a role for TLR3 in response to SARS-CoV-2^5^. Furthermore, pro-inflammatory cytokines can also be induced when PRRs detect damage-associated molecular patterns released during infection^36^. cGAS and STING have been shown to be activated in this manner by sensing of mitochondrial DNA released into the cytosol in Dengue virus-infected cells. Indeed, it has been proposed that the STING pathway mediates NF-κB activation and TNF induction in response to SARS-CoV-2^37^. In our work, loss of STING reduced but did not abolish the TNF response to SARS-CoV-2 (Fig 3B). However, it is important to note that some residual STING protein was still present in the targeted cells (Fig 3A), so it is possible that cytosolic DNA sensing by cGAS/STING induces the expression of TNF in response to SARS-CoV-2. Moreover, the study by Yin et al. reported that, in addition to MDA5-MAVS, the receptors LGP2 and NOD1 are also required for the type I IFN response to SARS-CoV-2 in Calu-3 cells^19^. LGP2 is an RLR and may function to amplify MDA5-mediated responses^38^, whereas NOD1 was recently found to play a role in facilitating RLR activation^39^. Canonically, NOD1 recognises bacterial peptidoglycans and signals *via* NF-κB to induce inflammatory cytokines^40^. In addition, peptidoglycan-free pathogens, such as viruses, as well as ER stress can lead to the activation of NOD proteins^41^. It is conceivable that NOD signalling and/or other pathways were responsible for the inflammatory cytokine response we observed in Calu-3 cells. Taken together, it is likely that the non-IFN pro-inflammatory cytokine responses we observed here resulted from the activation of PRRs other than MDA5.

Lastly, using flow cytometry, we observed upregulation of MxA protein, which is encoded by an ISG, in infected cell populations. This confirmed our RT-qPCR results showing induction of multiple ISG mRNAs upon infection. Staining for the viral nucleocapsid protein (N) allowed us to identify infected cells and uninfected bystander cells contained in the same sample. Interestingly, compared to infected cells, we observed more pronounced induction of MxA in bystander cells. This suggests that viral infection antagonised IFN receptor signalling. Indeed, other studies identified proteins in the IFNAR signalling cascade as targets of specific SARS-CoV-2 proteins. This includes the viral ORF6, which prevents STAT1 nuclear translocation and activation of ISG promoters^42,43^. Additionally, SARS-CoV-2 has been shown to downregulate protein expression of IFNAR1, JAK1 and TYK2, which are all involved in IFN receptor signalling^44^. Like SARS-CoV-2, SARS-CoV encodes viral proteins that interfere with the type I IFN system. For example, ORF3a from SARS-CoV targets IFNAR directly, and NSP1 inhibits STAT1 phosphorylation and IFNAR signalling by supressing host gene expression^45–47^. Therefore, it is likely that SARS-CoV-2 has additional proteins that block IFN receptor signalling^48^. Our work supports the idea that SARS-CoV-2 inhibits the response to type I IFNs, likely reducing host immunity and increasing virus propagation.

In sum, we show that SARS-CoV-2 infection activates MDA5, which – together with its downstream signalling partners MAVS and IRF3 – was essential for type I and III IFN production in response SARS-CoV-2 in Calu-3 cells. MDA5 was dispensable for the pro-inflammatory cytokine response that accompanied infection. In the future, it will be important to pinpoint molecular pathways that drive induction of pro-inflammatory responses, as they are highly relevant to COVID-19 disease progression and clinical outcome^49–53^.

## Materials and Methods

### Cell culture and virus infection

Cells were cultured at 37°C and 5% CO_2_ and routinely screened for mycoplasma contamination. Calu-3 cells (ATCC) were maintained in MEM (Gibco) supplemented with 10% v/v foetal calf serum (FCS, Gibco), 2 mM L-glutamine (Gibco), 1x sodium pyruvate (Gibco) and 1x non-essential amino acids (Gibco). Where indicated, cells were stimulated with IFN-A/D (R&D Systems) at 100 U/ml overnight. THP1 cells (kind gift from V Cerundolo) were maintained in RPMI (Sigma Aldrich) supplemented with 10% v/v (FCS) and 2 mM L-glutamine (Gibco). Where indicated, THP1 cells were treated with phorbol 12-myristate 13-acetate (PMA, 10 ng/ml; Invivogen) overnight prior to infection. All other cells (A549, kind gift from G Kochs; HEK293T, HEK293 and VERO E6, kind gifts from C Reis E Sousa; Huh7, kind gift from J McKeating) were maintained in DMEM (Sigma Aldrich) supplemented with 10% v/v FCS and 2 mM L-glutamine.

SARS-CoV-2 Victoria/02/2020 (passage 5) was produced by infecting Vero E6 cells at an MOI of 0.01 in DMEM supplemented with 1% FCS for 3 to 4 days. When cytopathic effects were visible, virus was harvested, aliquoted and frozen at −80°C. Virus stock was titrated by plaque assay. Briefly, virus was serially diluted ten-fold in DMEM with 1% FCS, and 100 μl of dilutions were added in quadruplicates to 24 well plates containing 2.5 x 10^5^ Vero E6 cells in 500 μl of medium. After incubation for 2 hours at 37°C, 500 μl of a semi-solid overlay (DMEM, 1% FCS, 3% carboxymethylcellulose) was added per well and the cells were incubated for 4 days. After removing the overlay and the medium, cells were washed in PBS, and fixed and stained using Amido Black stain for at least 30 minutes at room temperature. Plates were rinsed with water and dried before counting the plaques and calculating the titre.

To select the cell line to be used in our study, indicated cell lines were seeded in 25-cm^2^ flasks (1 x 10^6^ cells per flask) and infected the next day with SARS-CoV-2 at MOIs of 0.1 and 0.5. 2 hours post-infection, the medium was changed, and cells were returned to the incubator for 22 hours before analysis. For further experiments, cells were grown in 12-well plates (5.5 x 10^5^ cells per well) for 24 hours and infected with SARS-CoV-2 at an MOI 0.1 for 48 hours prior to processing.

### Lentiviral shRNA knockdown

Lentiviral plasmids encoding shRNAs targeting *GFP* (control; SHC005) and *MAVS* (06: TRCN0000149206; 45: TRCN0000148945) were obtained from the Sigma Mission library (Merck Darmstadt). Lentiviral particles for transduction were generated as follows: HEK293T cells were seeded in 6-well plates and the next day transfected with 1 μg psPAX2 packaging plasmid (Addgene 12260), 500 ng pMD2.G VSV-G envelope plasmid (Addgene 12259) and 1 μg shRNA plasmid using Fugene 6 (Promega). The next day, the medium was replaced. After another 24 hours, lentivirus-containing supernatant was harvested three times every 8-16 hours. Pooled supernatants were filtered through 0.45 μm filters and stored at −80°C. Calu-3 cells were transduced by addition of lentiviral supernatants containing 8 μg/ml polybrene (Merck Darmstadt). 48 hours after transduction, cells were selected by addition of media containing 5 μg/ml puromycin (Gibco). Surviving cells were used for experiments.

### CRISPR/Cas9 knockdown

The lentiCRISPRv2 plasmid backbone (Addgene #52961) was used for generation of bulk knockout cell populations^54^. Constructs targeting *GFP* (control, gctcgaactccacgccgttc), *MAVS* (ggccaccatctggattcctt), *IRF3* (ggtggtgcatatgttcccggg), *TMEM173* (STING) (gggtaccggagagtgtgctc), *DDX58* (RIG-I) (gaacaacaagggcccaatgg)^55^ and *IFIH1* (MDA5) (gtagcggaaattctcgtctg)^55^ were generated by restriction enzyme cloning using a published procedure^54^ with modifications. Complimentary oligonucleotides encoding the sgRNAs with suitable overhangs for ligation were annealed and phosphorylated using T4 Polynucleotide Kinase (NEB) with the T4 DNA Ligase Reaction buffer (NEB). The lentiCRISPRv2 vector was linearized by BmsBI (NEB) digestion, gel purified and ligated with prepared oligos using Instant Sticky-end Ligase Master Mix (NEB). After bacterial transformation, suitable clones were identified by Sanger sequencing. Lentiviral particle production and cell transduction was identical to shRNA lentiviruses but using pR8.91 packaging plasmid (kind gift from G Towers), pCMV-VSV-G (Addgene #8454) and Lipofectamine 2000 (Life Technologies).

### Immunoblotting

Cells were lysed with RIPA buffer (10 mM TRIS-HCl pH 8, 140 mM NaCl, 1% Triton-X100, 0.1% SDS, 0.1% sodium deoxycholate, 1 mM EDTA, 1 mM EGTA) and protein was quantified by BCA assay (Pierce). NuPAGE LDS sample loading buffer (Life Technologies) and 10% 2-mercaptoethanol were added to samples, which were then denatured by incubation at 95°C for 5 minutes. Samples were resolved by electrophoresis on 4-12% Bis-Tris gels with MOPS Running Buffer (Life Technologies NuPAGE system) and transferred to nitrocellulose membrane by electrophoresis at 100V for 1 hour in cold transfer buffer (Life Technologies) with 20% methanol. Membranes were blocked with 0.05% NP-40 (IGEPAL) in Tris-buffered saline (TBS-N; 50 mM NaCl, 50 mM Tris-HCl, pH 7.6) containing 5% non-fat milk (5% milk TBS-N) for 1 hour and probed with primary and HRP-conjugated secondary antibodies (GE Healthcare, 1:3000) diluted in 5% milk TBS-N for 1 hour at room temperature or overnight at 4°C, with rotation. Membranes were washed four times in TBS-N (5 minutes for each wash) after each antibody incubation. Proteins were visualised on an iBright (ThermoFisher) after exposure to Western LightningPlus-ECL chemiluminescent reagent (PerkinElmer). Primary antibodies included: beta-actin-HRP (AC-15, Sigma Aldrich), pSTAT1 (Y701) (58D6, Cell Signaling Technology), STAT1 (42H3, Cell Signaling Technology), pSTAT2 (D3P2P, Cell Signaling Technology), STAT2 (D9J7L, Cell Signaling Technology), hcGAS (D1D3G, Cell Signaling Technology), hSTING (D2P2F, Cell Signaling Technology), MyD88 (D80F5, Cell Signaling Technology), TBK1 (D1B4, Cell Signaling Technology), MAVS (ALX-210-929-C100, ENZO Life Science), IRF3 (D6I4C, Cell Signaling Technology), MDA5 (generated in house^55^) and RIG-I (clone ALME-I, ProSci #PSI-36-102).

### RT-qPCR

RNA was extracted from cells using RNeasy Plus Mini Kit (Qiagen, 74136) according to manufacturer’s instructions, quantified by Nanodrop, and stored at −80°C. RNA (1 μg) was converted into cDNA using SuperScript III Reverse Transcriptase (Thermo Fisher Scientific) and random hexamer primers (Qiagen, 79236) according to manufacturer’s instructions. cDNA was diluted to 100 ng/μl and quantitative PCR (qPCR) was performed using TaqMan Real-Time PCR Assays for designated genes and TaqMan Fast Advanced Master Mix (Thermo Fisher Scientific) according to manufacturer’s instructions. Assays were performed on QuantStudio 6 Flex Real-Time PCR machines (Thermo Fisher Scientific).

### Flow Cytometry

Cells were dislodged by washing in PBS and treatment with 0.05% trypsin (Gibco) for 20 minutes, followed by pipetting and centrifugation. For all subsequent steps, reagents were diluted in FACS buffer (PBS, 1% FCS, 2mM EDTA) unless stated otherwise, and washed twice with FACS buffer between steps. Cells were stained with Live/Dead Fixable Aqua Cell Stain (1:200 in PBS, Life Technologies, L34957) combined with FcR block (1:200 in PBS, eBioscience), fixed in 4% formaldehyde (10 minutes at room temperature), and permeabilised in PBS containing 0.1% Triton-X (20 minutes at room temperature). Cells were incubated with antibodies against human MxA (clone M143, kind gift from G Kochs) and SARS-CoV-2 N protein (clone EY-2A, kind gift from Alain Townsend^56^; 1:200, 30 minutes, 4°C), and goat anti-mouse AlexaFluor 488 (Life Technologies, A11029) and anti-human AlexaFluor 647 (1:500, 30 minutes, 4°C; Life Technologies, A21445), and resuspended in CellFix (1:10 in water; BD, 340181). Cells were analysed by flow cytometry on an Attune NxT Flow Cytometer (Thermo Fisher Scientific) and data were analysed using FlowJo software (BD).

## Acknowledgements

The authors thank William James for providing access to the BSL3 facility. The authors thank Arthur Huang, Pramila Rijal, Lisa Schimanski, Tiong Kit Tan and Alain Townsend for providing the anti-SARS-CoV-2 N antibody. This work was funded by the UK Medical Research Council [MRC core funding of the MRC Human Immunology Unit; JR] and the Wellcome Trust [grant number 100954; JR]. JH was supported by the European Commission under the Horizon2020 program H2020 MSCA-ITN GA 675278 EDGE. GF was supported by the Rothermere Fellowship and was also supported by Indspire’s Building Brighter Futures: Bursaries, Scholarships, and Awards program. RAR acknowledges the generous support of philanthropic donors that allowed funding from the University of Oxford’s COVID-19 Research Response Fund, which also supported the SARS-CoV-2 BSL3 facility. Initial funding for the Virus Screening Facility was provided by the Oxford BRC and Cancer Research UK. The funders had no role in study design, data collection and analysis, decision to publish, or preparation of the manuscript.

## Author contributions (using the CRediT taxonomy)

Conceptualization: NGS, LC, JH, AB and JR; Methodology: NGS, LC, JH and AB; Software: n.a.; Validation: NGS, LC, JH, AB and JR; Formal analysis: NGS, LC, JH, AB and JR; Investigation: NGS, LC, JH, AB, GF and JM; Resources: MD, RAR and MN; Data curation: NGS, LC, JH, AB and JR; Writing – Original Draft: NGS, LC, JH and JR; Writing – Review & Editing: all authors; Visualization: NGS, LC and JH; Supervision: JR; Project administration: JR; Funding acquisition: JR.

